# Ontology of RNA Sequencing (ORNASEQ) and its application

**DOI:** 10.1101/405720

**Authors:** Stephen A Fisher, Junhyong Kim

## Abstract

**Background:** Next-generation RNA sequencing is a rapidly developing technology with complex procedures encompassing different experimental modalities. As the technology evolves and its use expand, so does the need to capture the data provenance from these sequencing studies and the need to create new tools to manage and manipulate these provenance stores.

**Results:** Here we used the Ontology for Biomedical Investigations (OBI) and many other ontologies from the Open Biological and Biomedical Ontology (OBO) Foundry as a framework from which to create an application ontology (ORNASEQ: Ontology of RNA sequencing) to capture data provenance for next-generation RNA sequencing studies. Additionally, we provide an extensive real-life sample provenance data set for use in developing new provenance tools and additional sequencing data models.

**Conclusions:** The Ontology of RNA Sequencing (ORNASEQ) provides core terms for use in building data models to capture the provenance from next-generation RNA sequencing studies. The supplied sample provenance data also exemplifies many of the complexities of RNA sequencing studies and underscores the need for potent workflow management systems.

## Background

Until recently the cost of performing next-generation RNA sequencing (RNAseq) experiments limited the amount of data generated by a single lab and managing and properly documenting a few experiments was not fundamentally challenging.However, as sequencing costs have dropped, research groups are now running hundreds, thousands or even tens of thousands of RNAseq experiments, creating a need to systematically document experimental and informatics details and track provenance of the final published or publicly released datasets. RNAseq has also begun making its way into medical diagnostics, where data provenance is a necessity for quality assurance and regulatory compliance. Tracking the data provenance for hundreds or thousands of sequencing experiments in either a research or medical setting requires data models and structures that must be put into place to capture the necessary information at all stages of a sequencing experiment and it’s not always obvious what information is necessary. While there are numerous platforms and pipelines to analyze RNAseq data, there are limited data models or ontologies that could be applied to successfully capture the details of an RNAseq experiment [1–6].

Within a single next-generation sequencing experiment there is a dizzying amount of information that must be captured throughout the often-complex experimental procedures and post-sequencing informatics analyses. The problem is further complicated by the number of researchers or technicians who might be involved in a single sequencing experiment whose roles are interspersed in irregular patterns; complexity of biological specimens, their origin and experimental designs; and, the frequent disconnect between the biologists running the experiments and the bioinformaticists analyzing the data. Tracking data provenance spanning experiment procedures recorded in lab notebooks belonging to multiple biologists and computer log files residing in a series of cryptic directories on a file system quickly becomes an intractable problem. These challenges suggest a need for a comprehensive next generation sequencing provenance system. Data provenance requires data models, provenance models, and supporting infrastructure. Here, we focus on the first part of data models for RNA sequencing experiments and describe an Ontology for RNA Sequences (ORNASEQ). In addition, we provide a large next-generation sequencing use case from an active RNAseq workflow, using the PROV- XML database format for the community to use as an example dataset for development of provenance models and tools.

## Ontology for RNA sequencing

The Ontology for RNA Sequencing (ORNASEQ) is an application ontology based largely on the Ontology for Biomedical Investigations (OBI)[1], using the principles of OBO Foundry[7]. Specifically, ORNASEQ contains 162 terms, 117 of the terms are from 16 existing ontologies, with 28 new terms having now been added to OBI and 17 terms being defined directly in ORNASEQ (see Table 1). ORNASEQ is designed to annotate RNA-based next-generation sequencing, although much of ORNASEQ would also apply to DNA-based next-generation sequencing. The ontology was designed, in part, through efforts to track the data provenance of thousands of RNAseq samples collected by the NIH Common Fund Single Cell Analysis Program- Transcriptome (SCAP-T) program[8]. Terms included in the ontology cover pre-sequencing preparations and primary post-sequencing data analysis.

**Table 1:** The set of external ontologies included in ORNASEQ.

| Ontology | Number Terms |
| --- | --- |
| **BFO:** Basic Formal Ontology<br>http://purl.obolibrary.org/obo/bfo.owl | 10 |
| **CHEBI:** Chemical Entities of Biological Interest<br>http://purl.obolibrary.org/obo/chebi.owl | 2 |
| **CL:** Cell Ontology<br>http://purl.obolibrary.org/obo/cl.owl | 6 |
| **EFO:** Experimental Factor Ontology<br>http://www.ebi.ac.uk/efo/efo.owl | 6 |
| **GENEPIO:** Genomic Epidemiology Ontology<br>http://purl.obolibrary.org/obo/genepio.owl | 4 |
| **GO:** Gene Ontology<br>http://purl.obolibrary.org/obo/go.owl | 5 |
| **IAO:** Information Artifact Ontology<br>http://purl.obolibrary.org/obo/iao.owl | 14 |
| **NCBITaxon:** NCBI Organismal Classification<br>http://purl.obolibrary.org/obo/ncbitaxon.owl | 1 |
| **NCIT:** NCI Thesaurus OBO Edition<br>http://purl.obolibrary.org/obo/ncit.owl | 6 |
| **OBI:** Ontology for Biomedical Investigations<br>http://purl.obolibrary.org/obo/obi.owl | 79 |
| **OBIws:** OBI web service, development version<br>*http://purl.obolibrary.org/obo/obi/webService.owl* | 5 |
| **OGMS:** Ontology for General Medical Science<br>*http://purl.obolibrary.org/obo/ogms.owl* | 1 |
| **OMIABIS:** Ontologized MIABIS<br>*http://purl.obolibrary.org/obo/omiabis.owl* | 2 |
| **SO:** Sequence Types and Features<br>*http://purl.obolibrary.org/obo/so.owl* | 1 |
| **TAXRANK:** Taxonomic rank vocabulary<br>*http://purl.obolibrary.org/obo/taxrank.owl* | 2 |
| **UBERON:** Uberon multi-species anatomy ontology<br>*http://purl.obolibrary.org/obo/uberon.owl* | 1 |

### Pre-Sequencing Provenance

Knowing what happens to a data sample prior to sequencing is essential to understanding the analyzed data. The provenance surrounding the preparation of sequencing data can prove invaluable to diagnosing aberrant results. These protocols are often revised over time and as hardware and reagents evolve and a multi-year study will likely include many versions of protocols with varying degrees of differential change. Even in the controlled context of medical diagnostics, changes are inevitable, for example, when a reagent becomes discontinued or hardware are upgraded.

When capturing experimental lab data provenance, it is difficult to know what to capture and in what format (e.g. as fields in a database, as a Word or PDF document, or even as a reference to an entry in an electronic notebook). As sequencing preparation protocols are often distributed as PDF or Word documents, trying to track changes across multiple such documents quickly becomes a tedious process that is difficult to automate and nearly impossible to query for specific questions (e.g., what version of sequencing chemistry was used for what samples). Conversely, because the protocols involve many incremental steps and details that are inter- related in a complicated manner, it is difficult to convert each protocol into a fine-grained structured knowledge model suitable for standard DBMS. Similar to the scheme implemented by the Genomic Standards Consortium [3], we propose experimental provenance be captured in a hybrid fashion including both detailed protocol files and hardcoded fields tracking fundamental datum external to the protocol files. In this RNAseq data provenance schema, complete protocol definitions are stored in the provenance database as files (typically PDF or Word documents). Critical or commonly changed features of the protocol are additionally captured in the database schema. While not optimal, this approach preserves the fine detail required to replicate an experimental procedure, while allowing for structured query of the main features.

### Post-Sequencing Provenance

When RNAseq data comes off the sequencer it is typically converted into fastq files by a proprietary, sequencer-specific program. A fastq file is a text file containing nucleotide sequences [9]. In the case of RNAseq, fastq files contain nucleotide sequences representing the RNA molecules from a biological sample. Any one RNA molecule might be represented by tens to hundreds of thousands of sometimes duplicate nucleotide sequences in the fastq file, with each nucleotide sequence referred to as a “read” (i.e. a read out of the biological sequence). The fastq files are run through what is called a “primary analysis pipeline”, which may include any number of steps such as generating quality control metrics, removing contaminant sequences from one or both ends of the reads, removing low quality sub-sequences from reads, removing duplicate reads, and aligning reads to a pre-defined reference library. Primary analysis pipelines and more generally RNA sequencing experiments often culminate in the generation of gene-, transcript-, or exonic-counts of the number of reads associated with each of these categories. The various steps will mostly remain consistent within a research group, a project or a medical diagnostic test. However, programs evolve, algorithmic bugs are fixed, and reference libraries are refined. As with wet lab procedures, it is common and often necessary for primary analysis pipelines to change by varying degrees over time. Since the data and processes for the informatics steps are already machine readable, capturing provenance for RNAseq analysis pipeline is a more obvious task than in a wet lab context. However, relevant provenance information is typically realized as a set of log files spread across a series of programs, each with specific directory structures. Using log files to track provenance data also quickly breaks down as programs change or pipelines evolve and often requires complex programming to perform simple queries across data samples. As with tracking wet lab data, provenance from sample processing pipelines needs to be rigorously captured and systematically stored in a central database. Addressing these challenges require incorporation of a well-defined and use-case oriented ontology for the provenance objects, which we provide with ORNASEQ.

## Sample Data

ORNASEQ is meant to provide core terms used to track the provenance of RNAseq datasets. However, any particular experiment or RNAseq use case will require a multitude of additional terms. Here we provide a dataset containing curated and modified data provenance from 1,347 next-generation sequencing samples. The dataset contains 93 data fields, using ORNASEQ terms as appropriate. Each sample was processed with one of four different versions of a primary analysis pipeline and there was from one to three variants (sub-versions) of each pipeline version. Specifically, samples were processed either as “single-end” or “paired-end” and aligned with either STAR[10] or Bowtie[11, 12], ultimately leading to nine possible pipeline variants. The dataset is provided as PROV-XML (see Additional file 2), with the data summarized in an Excel table (see Additional file 1). Real world use cases usually include messy data. Occasionally pipelines are run incorrectly either through intentional operator actions or error. It’s also quite common to have missing or incomplete data provenance again through user actions (e.g. data was pulled from a source that lacked sufficient provenance), programming errors, computer issues, etc. The dataset provided here intentionally includes both incorrect and missing data.

### Primary Analysis Pipeline Stages

The data provided describes an analysis pipeline with seven possible stages but with each analysis only including five of the seven stages. Table 2 includes a subset of data tracked for each pipeline stage and which stages might be used in a particular pipeline version. For example, the stage HTSeq was only used in pipeline version 1.0 while VERSE was only used in later pipeline versions. The stages are very briefly described here and more complete descriptions can be found in the PennSCAP-T Pipeline[13].

**Table 2:** Subset of pipeline-specific parameters included in sample dataset, across pipeline stages and pipeline versions.

| STAGE | PARAMETER | VALUE |  |  |  |
| --- | --- | --- | --- | --- | --- |
| **BLAST** | blastn version | 2.2.25 | 2.2.30 | 2.2.30 | 2.2.30 |
|  | parseBlast.py version | 1.1 | 1.5 | 1.5 | 1.8 |
| **FASTQC** | fastqc version | 0.10.1 | 0.11.2 | 0.11.2 | 0.11.2 |
| **TRIM** | trimReads.py version | 0.6 | 0.6 | 0.6 | 0.7.2 |
|  | removeN | 1 | 1 | 1 | 1 |
|  | minLen | 30 | 20 | 20 | 20 |
|  | phredThresh | - | 53 | 53 | 53 |
|  | numAT | 30 | 26 | 26 | 26 |
| **BOWTIE** | bowtie version | 0.12.7 | 1.1.1 |  | 2.2.9 |
|  | minins | 250 | 150 |  | 150 |
|  | maxins | 450 | 600 |  | 600 |
| **STAR** | star version | 2.3.0.1 | 2.4.0h1 | 2.4.0j | 2.5.1b |
|  | samtools version | 1.1 | 1.1 | 1.2 | 1.2 |
| **HTSEQ** | htseq version | 0.5.4p5 |  |  |  |
| **VERSE** | verse version |  | v0.1.4 | v0.1.4 | v0.1.5 |
| **Pipeline Version** |  | **1.0** | **2.0** | **2.0.1** | **2.1** |
| **Number Samples** |  | **208** | **413** | **216** | **510** |

- BLAST - a subset of samples was aligned to the Blast NR database, using BLAST as a quality control check.
- FastQC – FastQC is used as an additional quality contro check.
- TRIM – contaminant sequences were removed from reads.
- BOWTIE – Bowtie was used to align reads to a reference genome.
- STAR – STAR was used to align reads to a reference genome.
- HTseq – HTSeq was used to assign aligned reads to genes.
- VERSE – VERSE was used to assign aligned reads to genes.

In the accompanying dataset, samples were aligned with either Bowtie or STAR but not both. Similarly, alignments were processed by either HTSeq or VERSE, but not both. The Excel table highlights which provenance terms are consistent within a pipeline version. For example “BLAST.blastn.version” has pipeline version-specific values. Provenance terms that might contain incorrect values are also denoted. For example, the value for “STAR.star.version” in Pipeline version 2.0.1 might be missing.

## Conclusion

Next generation sequencing (NGS) is a complex technology with multiple varied steps. Capturing the provenance of NGS data requires complex systems and multi-user collaborations. The schemas and ontology we define here offer a basic framework that can be tuned and expended by researchers to the particulars of individual studies, while providing basic commonalities across studies.

The dataset provided illustrates some of the complexities of the data provenance from downstream processing of NGS data. These complexities will grow as the field evolves. For example, there are now hundreds of variants of basic sequencing protocol that are specific to particular biology applications (e.g., ATAC-seq; [14]; Drop-Seq;[15, 16]). Each of these involve variations in experimental and informatics processes. It will become necessary to build workflow management systems and smart clustering algorithms of the provenance[17] to help segment incomplete and erroneous data. We propose that our large real-life dataset example will prove useful in designing future new workflow systems and provenance models.

## List of Abbreviations

### **OBI:** Ontology for Biomedical Investigations

### **ORNASEQ:** Ontology of RNA Sequencing

### **OBO:** Open Biological and Biomedical Ontology

### **RNAseq:** next-generation RNA sequencing

### **SCAP-T:** NIH Common Fund Single Cell Analysis Program-Transcriptome

### **NGS:** Next generation sequencing

## Declarations

### Ethics approval and consent to participate

not applicable

### Consent for publication

not applicable

### Availability of data and material

The ontology presented in the current study is available from GitHub, http://doi.org/10.5281/zenodo.1311869. The sample dataset presented in the current study is available as additional article files.

### Competing interests

The authors declare that they have no competing interests.

### Funding

This work was supported in part by NIH grants 5U01EB020954 and U01MH098953. The NIH played no role in the results of this research effort or the text of this paper.

### Authors’ contributions

SF and JK contributed to the design of this study and preparation of the manuscript.

## Acknowledgements

We are grateful to Christian Stoeckert and Daniel Berrios for valuable feedback and assistance adding terms to the Ontology for Biomedical Investigations.

## Additional Files

### Additional file 1

### **Format:** Excel Workbook from Excel for Mac version 16 (.xlsx)

### **Title**: A Large Curated RNAseq Metadata Dataset

### Description

This file contains a curated dataset consisting of metadata collected from 1347 next-generation sequencing samples. The file also contains a set of terms that can be used to separate the data samples between nine primary analysis pipelines

## Additional file 2

### **Format**: XML file (.xml)

### **Title**: PROV-XML Instantiation of a RNAseq Metadata Dataset

### Description

This is an XML file that contains a PROV-XML representation of the data included in “Additional file 1.xslx” that is, an example dataset containing metadata from 1347 RNAseq data samples that were processed with one of nine primary analysis pipelines.

